# Social buffering as an indirect effect: mixed-effects modeling approaches

**DOI:** 10.1101/2025.01.15.633205

**Authors:** M.A. Sekhar, N.A. Dochtermann

## Abstract

1. The potential for an individual’s social partners to buffer—or otherwise modify— how individuals respond to their environment has been demonstrated to be important in many contexts. This buffering has the potential to affect responses to human modifications of environments. Unfortunately, statistical tools for identifying buffering effects have not been well developed.
2. Here, we demonstrate how social buffering fits into the context of a phenotypic equation conceptual approach and then connects to mixed-effects modeling for estimating buffering and other modifying effects of social behavior.
3. We explore the power and accuracy for buffering in response to known environments, providing a guide for empirical investigation. We found that increasing the sampling of social interactions decreases bias and increases precision and power to a greater extent than increasing sampling of focal individuals.
4. We also introduce how buffering in response to unknown environmental variation can be statistically modeled and tested. If environments are unknown, social buffering can be statistically tested for using double hierarchical generalized linear models by including social partner identity as a random effect that influences residual variation.
5. Finally, we discuss how these approaches fit into the broader literature on indirect effects and indirect genetic effects. Placing social buffering in the context of indirect effects reveals that the evolution of social buffering is affected by both variation in an individual’s behavior and variation in how individuals affect each other. This has important implications for social and evolutionary organismal responses to changing environments.

## Introduction

Organisms regularly experience changing abiotic and biotic environmental conditions that require phenotypic responses and modifications. These environmental changes can happen suddenly, seasonally, or on longer scales and include temperature fluctuations, extreme weather events, shifts in climate patterns, and other effects (Lawson et al. 2015). Due to anthropogenic influences, environmental variability is also expected to increase (IPCC 2023).

These organismal responses to environmental variability are via phenotypic plasticity: differential phenotypic expression by the same genotype. Phenotypic plasticity is well documented for physiology, morphology, life-history, and behavior (e.g. Noble et al. 2018, Stamp and Hadfield 2020, MacLeod et al. 2022). In many organisms, this plasticity can also be induced by an individual’s social partners and neighbors (Moore et al. 1997, Wolf et al. 1998, Wolf et al. 1999).

When partners or neighbors induce plasticity that improves a focal individual’s fitness, decreases negative effects of environmental variation, or otherwise benefits the focal individual, this plasticity has been termed “social buffering” (sensu Kikusui et al. 2006). For example, in Barbary macaques (*Macaca sylvanus*), both low temperatures and aggression from conspecifics results in physiological stress as measured by glucocorticoid metabolite levels (Young et al. 2014). However, individual macaques that had stronger social bonds— as measured by allogrooming, contact, and other behaviors—showed decreased glucocorticoid levels, indicating decreased stress and a buffering of harmful effects by social partners (Young et al. 2014).

Defining social buffering as plasticity modified by social partners allows for a clear conceptual and statistical framework that can be used to improve our understanding of its biological effects. Specifically, the evolutionary ecological concepts of the phenotypic equation (Nussey et al. 2005, Nussey et al. 2007, Westneat et al. 2015) and of indirect effects (Moore et al. 1997, Wolf et al. 1998, Wolf et al. 1999, Bailey et al. 2018) can be used to understand social buffering. Both concepts are readily translated to statistical models, facilitating rigorous research into buffering. Here, we outline this conceptual framework and evaluate how well its statistical application identifies variation among individuals in buffering effects.

### A conceptual and statistical framework for social buffering

If we define social buffering as phenotypic plasticity, we can build a conceptual framework for its understanding starting from simple models.

First, we can define plasticity as a response to an abiotic or biotic environmental variable. Following Nussey et al. (2007), this can be represented as a “phenotypic equation” (Figure 1A). The phenotypic equation approach involves translating a biological hypothesis about the causal influences on phenotypes into a mathematical expression. At the simplest level, the phenotypic equation represents the phenotype of interest and plasticity in a simple linear model where the phenotype of interest (y) is a function of a single environmental variable (*x*):

**Figure 1.**
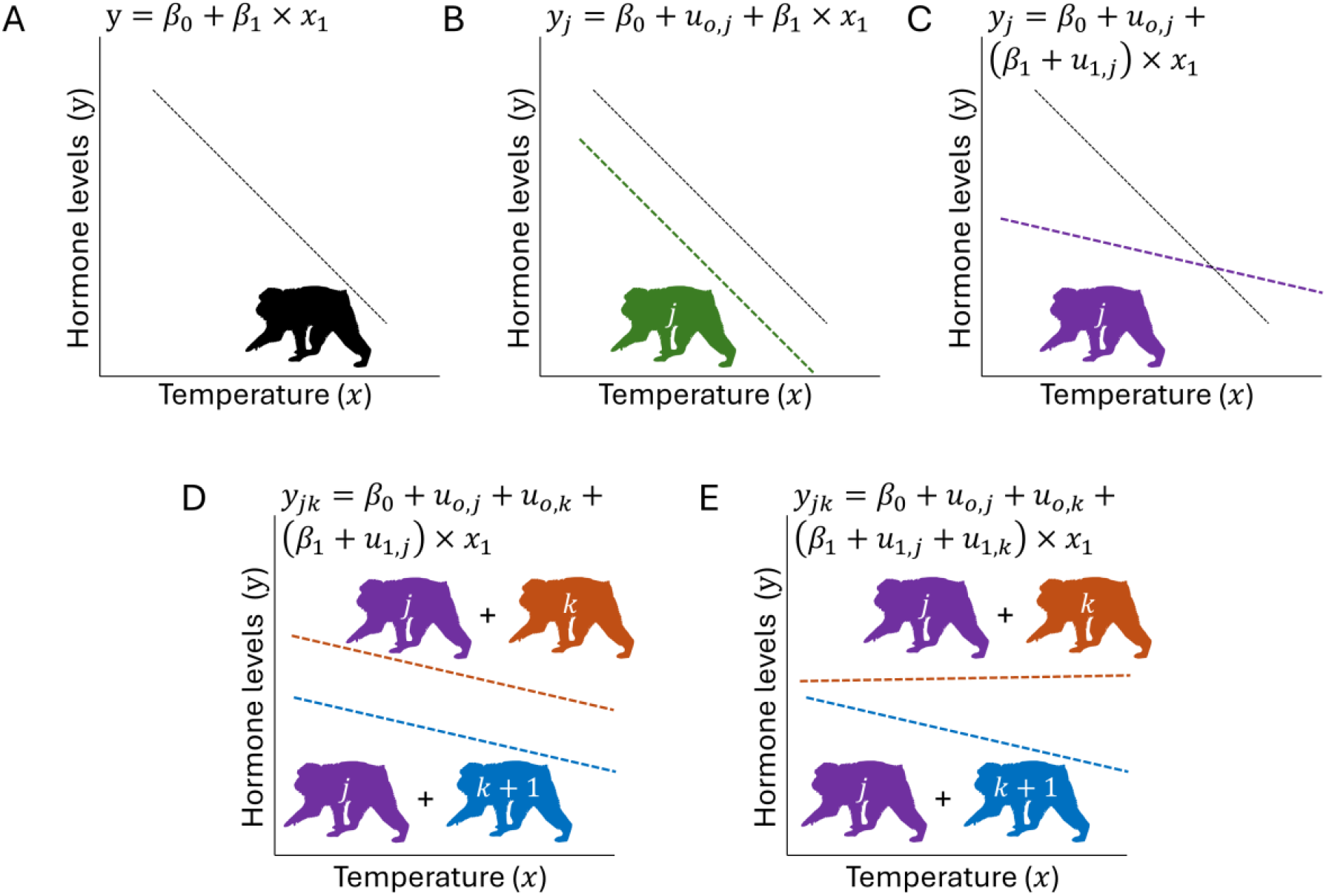
Social buffering as induced plasticity. A) A simple model of plasticity where a response variable y (e.g. hormone levels in macaques (Young et al. 2014) is influenced by some environmental variable, x (e.g. temperature). B) An individual, j, might differ in their average hormone levels (by µ_0,j_) or their average hormone level and the degree of plasticity they express (C, by µ_1,j_). D) A conspecific, k, might influence the average hormone levels expressed by j (by µ_0,k_). E) With social buffering, conspecifics (k and k+1) differentially influence the degree of plasticity j expresses (by µ_1,k_). Macaque silhouettes were produced by K.R. Caspar under creative commons license CC BY 3.0 (link) via phylopic.org and recolored by the authors.

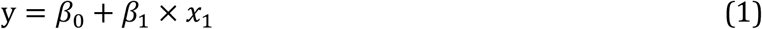

This models population-level plasticity where the average phenotype at *x*= 0 is represented by β_*0*_ and the magnitude of plasticity is given by β_1_. β_1_ is interpreted as the amount by which y increases for a single unit increase in *x*. In our prior example, this captures the biological response of macaques where decreasing temperatures result in increasing hormone levels (Figure 1A). This model of plasticity is simply fit with linear regression.

Before introducing social buffering, we can first expand the phenotypic equation to capture the fact that individuals often differ in their average phenotypes (Bell et al. 2009). We do so by adding a term that captures an individual’s average deviation from the population-level model of plasticity:

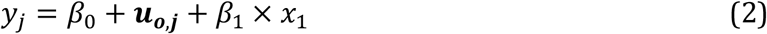

where *y*_*j*_represents the phenotype of individual “*j*” at a particular value of x. *u*_*o,j*_is the degree to which individual j deviates, on average, from the population level plastic response given by equation 1. For example, macaque j could, on average, have lower hormone levels than other macaques (Figure 1B). However, in this formulation, the degree of plasticity does not differ among individuals. This expanded phenotypic equation can be fit using mixed-effects models when there are repeated measurements per individual by including random intercepts for individuals (Dingemanse and Dochtermann 2013).

We can further expand the phenotypic equation by recognizing that not only might individuals differ in their average behavior, but also in their response to environmental conditions (i.e. phenotypic plasticity; Nussey et al. 2007, Dingemanse et al. 2010, Dingemanse and Dochtermann 2013, Harrison et al. 2018). As in equation 2, this can be represented by adding a term for the individual’s deviation. Now the individual is deviating from the population-level slope and intercept (Figure 1C):

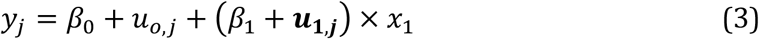

*u*_1,*j*_is the degree to which individual j deviates from the average slope (β_1_). Biologically, this would represent macaque j showing a different hormonal response to changing temperatures than do other macaques. To fit this conceptual model to data, repeated measurements per individual across a range of values of x are needed. With such data, the modeled response can be fitted using a mixed-effects model with both random intercepts and random slopes.

With equation 3 representing our conceptual model of plasticity for focal individual j, we can now add the effects of social partners. While there are several ways to conceptualize this, the simplest is to recognize that a social partner, e.g. macaque k, may affect our focal individual’s average behavior (Figure 1D):

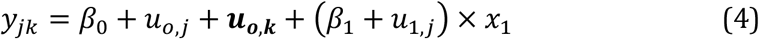

and the focal individual’s plasticity (Figure 1E):

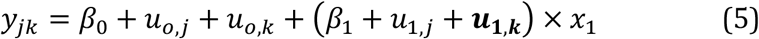

where *u*_*o,k*_is individual k’s effect on j’s average behavior and *u*_1,*k*_is k’s affect on the plastic response.

Importantly, equation 4 and the term *u*_*o,k*_do not capture social buffering. While *u*_*o,k*_is plasticity in response to a social partner, it does not represent a change in the plastic response to x. Instead, given our earlier definition, equation 5 and *u*_1,*k*_includes social buffering. This model of social buffering can be fit using mixed-models where random intercepts and random slopes are fit to social partner identity. To fit equation 5 and estimate variation in social buffering, an individual would have to have its phenotype measured repeatedly over a range of values of x and with multiple social partners.

Representing social buffering according to equation 5 and fitting its effects as a random slope model changes how buffering is considered. Previous empirical work on social buffering has focused on how social partners alter the mean phenotype of focal individuals, equation 5 emphasizes variance in these effects. For example, in a classic example of social buffering, Coe et al. (1982) showed that vocalizations and activity of squirrel monkeys (*Saimiri sciureus*) in response to a snake decreased when focal individuals were with social partners. In contrast, the approach represented by equation 5 would identify whether social partners differentially affect a focal individual’s behavior. This would be identified as variance in *u*_1,*k*_. In the case of Coe et al.’s (1982) squirrel monkeys, representing buffering as in equation 5 would allow testing of whether social partners differ in how they affect focal individual vocalizations and partner activity.

Equations 4 and 5 extend Nussey et al.’s (2007) phenotypic equation to include the indirect effects of social partners. Indirect effects refer to the influence of a social partner on the phenotype of a focal individual (Moore et al. 1997, Wolf et al. 1999). Social buffering thus represents a particular type of indirect effect. Specifically, social buffering is an indirect effect on the plasticity of a focal individual by social partners. If relatedness among individuals is known, either through pedigree information or genomic relatedness, equation 5 and social buffering can also be extended to separately estimate how the genotype of a partner contributes to social buffering using animal models (Kruuk 2004).

Placing social buffering in the context of the phenotypic equation provides a clear conceptual and statistical framework for its study. Relatedly, placing social buffering in the context of indirect effects identifies important evolutionary consequences. For example, as an indirect effect, social buffering can have profound effects on the evolution of plasticity and the ability of populations to evolve in response to changing environments (Moore et al. 1997, Santostefano et al. 2024). We explore these evolutionary implications further in the discussion.

Regardless, with social buffering modeled as in equation 5, we can now ask whether buffering can be identified statistically.

### Mixed-effects modeling of social buffering

Among-individual variance in parameter *u*_1,*k*_in equation 5 can be estimated in a mixed-effects modeling framework but how well?

To address this we conducted a bias, precision, and power analysis. Bias estimates systematic differences between the estimated and “true” values, indicating over-or underestimation. Precision reflects the consistency of estimates. Power represents the ability to detect effects, depending on sampling and effect magnitude (Cohen 1988). Combined, these measures tell us how well social buffering can be estimated and the conditions under which it can be statistically detected.

We simulated social buffering as in equation 5 but with additional random deviations (i.e. residuals) added to each (i) phenotypic expression (e_ijk_). All focal (*u*_*0,j*_and *u*_1, *j*_) and social partner effects (*u*_*0,k*_and *u*_1,*k*_) were modeled as independent variables. However, in natural populations, these effects may not be entirely independent. For demonstration purposes, we assumed that the direct and indirect effects on plasticity (i.e. *u*_1,*j*_and *u*_1,*k*_) were of equal magnitude and set to one-tenth the magnitude of effects on average responses (Table 1).

**Table 1.** Simulation parameters corresponding to equation 5 but with additional random deviations (residual variation). The “Value” column lists the distributional descriptions or specific value used in bias, precision, and power simulations. “Variance Components” are the equations for the contribution of a particular parameter to the total population variance in y (see Allegue et al. 2017). “Variance Contribution” lists the realized variances (using the specific values in the component equations).

| Parameter | Parameter Type | Value | Variance Component | Variance Contribution | Variance Proportion |
| --- | --- | --- | --- | --- | --- |
| $\beta_0$ | Population-level intercept | 0 | 0 | 0 | 0 |
| $\beta_1$ | Population-level slope | 1 | 0 | 0 | 0 |
| $x_1$ | Independent variable | $N(0,0.5)$ | $\beta_1^2 Var(x_1)$ | 0.5 | $\sim 0.21$ |
| $\mu_{0,j}$ | Focal Individual-level intercept | $N(0,0.5)$ | $Var(\mu_{0,j})$ | 0.5 | $\sim 0.21$ |
| $\mu_{1,j}$ | Focal Individual-level slope | $N(0,0.1)$ | $Var(\mu_{1,j})Var(x_1) + E(x_1)^2 Var(\mu_{1,j})$ | 0.05<sup>a</sup> | $\sim 0.02$ |
| $\mu_{0,k}$ | Partner-level intercept | $N(0,0.5)$ | $Var(\mu_{0,k})$ | 0.5 | $\sim 0.21$ |
| $\mu_{1,k}$ | Partner-level slope | $N(0,0.1)$ | $Var(\mu_{1,k})Var(x_1) + E(x_1)^2 Var(\mu_{1,k})$ | 0.05<sup>a</sup> | $\sim 0.02$ |
| $e_{i,j,k}$ | Random deviation (residual) | $N(0,0.8)$ | $Var(e)$ | 0.8 | $\sim 0.33$ |
| Total variance |  |  |  | 2.4 | 1.0 |

<sup>a</sup> because $x_1$ is modeled as having an average of 0, its contribution ( $E(x_1)^2$ ) cancels out

Using the population parameter values listed in Table 1, we simulated population samples across 54 different population and sampling schemes. These sampling schemes varied based on number of focal individuals, number of social partners each focal individual interacted with, and the number of times the focal individual interacted with each partner. We generated samples of 10, 25, 50, 100, 150, and 200 individuals. These individuals interacted with 3, 5, or 10 social partners. Focal individuals then interacted with each social partner 3, 5, or 10 times. For the lowest amount of sampling, this generated sample sizes of 90: 10 focal individuals interacting with 3 social partners and doing so 3 times with each social partner. For the greatest amount of sampling this generated sample sizes of 20,000. To these simulated data sets, we fit random intercept or random intercept plus random slope models corresponding to equations 4 or 5. Each combination of sample sizes was fit to 1000 independent samples.

After fitting these 54,000 sample sets to equations 4 and 5, we estimated bias, precision, and power. Bias and precision were estimated for the variance among social partners in the plasticity they induced in partners (i.e. V ar(μ_1,*k*_)). Bias was calculated as the mean difference between the estimated value and the simulated value (Table 1). We also calculated the standard error of this difference. Precision was calculated as the root mean squared error (RMSE):

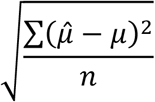

While RMSE has a defined standard error (Faber 1999), this standard error assumes that deviations are normally distributed. Rather than making this assumption, we calculated uncertainty in RMSE by bootstrapping 95% confidence intervals. Finally, because lower values of RMSE indicate more precise estimates, this value can more directly be interpreted as imprecision.

To determine power to detect social buffering, for each sample set we conducted a likelihood ratio test comparing the fit of mixed effects models corresponding to equations 4 and 5. We then recorded whether the addition of the random slope term for the social partners significantly improved fit.

All simulations were conducted in the R statistical language (v4.4.1; R Core Team 2024) and models were fit using the <monospace>glmmTMB</monospace> library (Brooks et al. 2017).

Because we investigated a narrow range of sampling schemes and a single set of parameters, we have also provided an R function for bias, precision, and power analysis that can be used by researchers for their own prospective power analyses (https://osf.io/tg39c).

### bias, precision, and power in estimating social buffering via mixed-effects modeling

As expected, for the parameter values explored here, bias decreases while precision and power increase with increasing sampling. However, how this effort is allocated---whether by measuring responses of more focal individuals, responses of more social partners, or multiple responses to each social partner—substantially impacts the estimation and detection of social buffering.

There was a slight bias toward zero in the estimate of the variance in social buffering (_V ar(μ1,*k*_)), particularly when only 10 focal individuals were sampled and sampled infrequently (Figure 2A – C). This bias approached zero as the number of social partners and the number of times a focal individual interacted with these partners increased (Figure 2A – C). Importantly, this bias was rarely more than 10% of the actual parameter. However, the smallest sample sizes were an exception to this and in these cases variance in social buffering was often estimated as approaching zero or as zero. Similarly, precision increased as sampling increased (Figure 2D – F). Bias and imprecision decreased more quickly with more social partners and interactions per partner versus an equivalent increase in the number of focal individuals.

**Figure 2.**
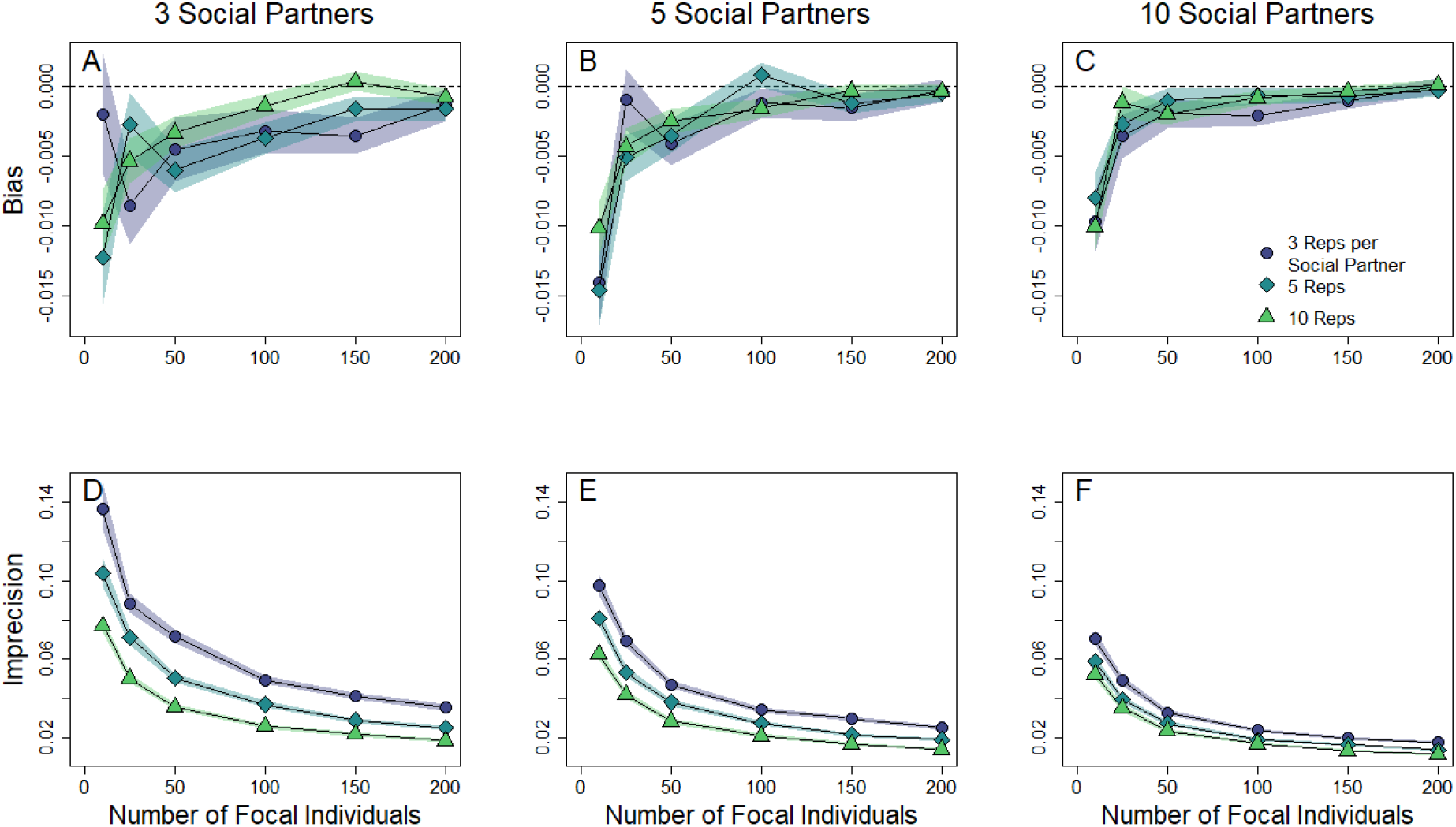
Bias (A – C) and precision (D – F) of estimates of variance in social partner induced plasticity (µ_1,k_, equation 5, Figure 1). Bias and precision were estimated across a range of sample sizes in focal individuals (x-axis). Focal individuals were simulated as interacting with 3 (A & D), 5 (B & E) or 10 (C & F) social partners. Focal individuals were simulated as interacting with either 3 (panels A & D), 5 (panels B & E), or 10 (panels C & F) social partners. For each scenario, focal individuals interacted with each social partner multiple times, represented by three levels of repetitions: 3 times (circles), 5 times (diamonds), or 10 times (triangles). These repetitions are shown as separate lines in each panel, with the shapes corresponding to the respective number of interactions per social partner. Each point represents 1000 simulations. A – C) Bias was estimated as the difference between estimated values and the simulated population values (i.e. 0.1). At the smallest sample sizes of focal individuals, variance in µ_1,k_ was underestimated by around 10 – 15%. Shaded portions cover the standard errors of estimated bias. D – F) Imprecision (lower values indicating greater precision) was estimated as the root mean square error and, as expected, decreases with increasing sampling. Shaded portions cover bootstrapped 95% confidence intervals. Total sample sizes are the product of the number of focal individuals, the number of social partners, and the number of replications per social partner.

Power also generally increased rapidly with sampling (Figure 3). As was the case for bias and precision, the increase in power was strongly dependent on the number of social partners and number of interactions with these partners. When individuals interacted with social partners infrequently (3 times) and with few social partners (3), power only exceeded 0.8 with 150 focal individuals (Figure 3A).

**Figure 3.**
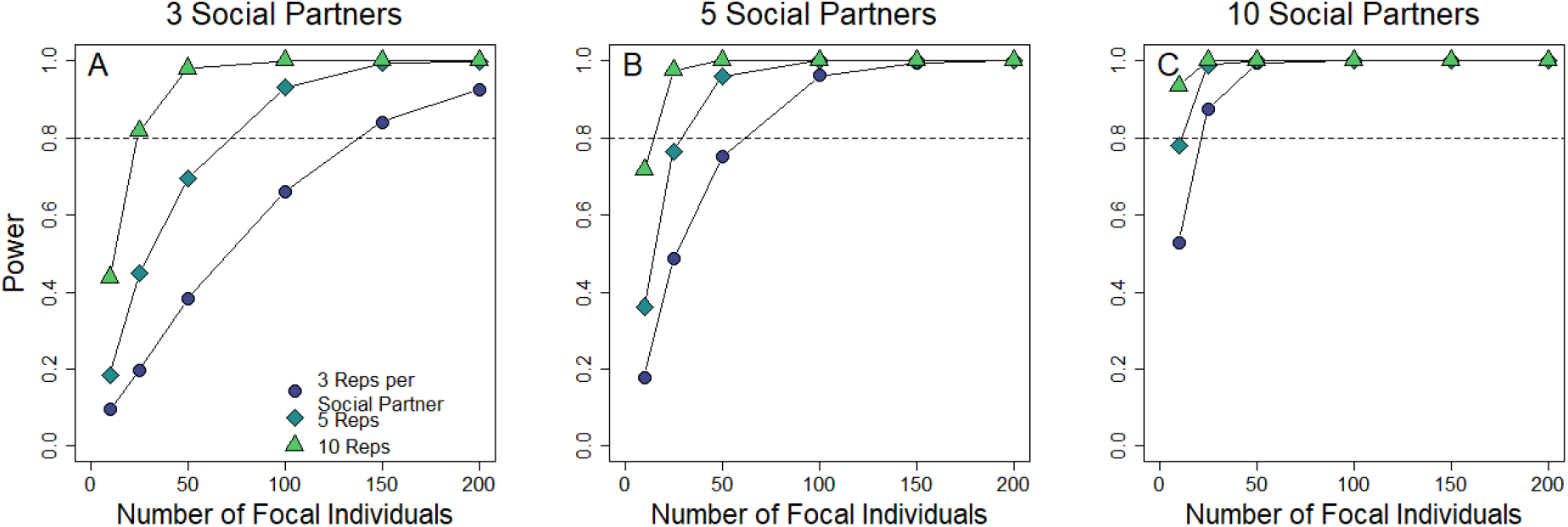
Statistical power to detect variance in social buffering. “Significant” social buffering was tested via a likelihood ratio test of the inclusion/exclusion of social partner induced changes in slope (µ_1,k_, equation 5, Figure 1) with α = 0.05. The proportion of 1000 simulations where inclusion of µ_1,k_ significantly improved model fit was calculated for a range of sample sizes in focal individuals interacting with 3 (A & D), 5 (B & E) or 10 (C & F) social partners. Each focal individual interacted with a social partner multiple times: 3 (circle), 5 (diamond) or 10 times (triangle). The dashed line indicates a power threshold of 0.8. Total sample sizes are the product of the number of focal individuals, the number of social partners, and the number of replications per social partner.

Put another way, increasing the number of social partners and interactions with these social partners led to a greater increase in power versus a similar increase in sampling that emphasized increasing the number of focal individuals sampled (Figure 3). For example, power when sampling 10 focal individuals interacting with 10 social partners 10 times each (total sample size = 1000) was 0.94. In contrast, sampling 100 focal individuals interacting 3 times with 3 partners (total sample size = 900) provided a power of 0.66.

While the required sample sizes to reliably identify social buffering are large, they represent sample sizes increasingly used in studies of social behavior (e.g. Smith et al. 2018, Ogino et al. 2023). For example, social behavior research with vampire bats (*Desmodus rotundus*) often includes many thousands of observations (Hartman et al. 2024). Moreover, our simulations identify that priority should be given to increased sampling of fewer focal individuals, rather than shallower sampling of more individuals. The required sample sizes are also likely common but obscured via summarization before analysis: a mixed-effects approach does not require averaging at the focal or partner level as each data point is not assumed to be independent of others.

The use of mixed-effects models to test for social buffering also allows for more appropriate examination of connections between buffering and fitness. Selection is typically underestimated for labile, repeatedly expressed traits (Dingemanse et al. 2021), like those affected by social partners. This underestimation comes about because fitness is often calculated based on a single phenotypic measure or the average of multiple measurements. Correlations between means misestimate relationships between phenotypic traits (Dingemanse et al. 2012) or, here, between socially induced plasticity and fitness. The mixed-effect model of social buffering given in equation 5 can be combined with “error-in-variable” and “social animal model” approaches (Ponzi et al. 2018, Martin and Jaeggi 2022) to properly estimate the relationship between social buffering and fitness.

### Social buffering in response to unknown environmental variables

Known variables allow for direct modeling by including covariates and corresponding random slope terms. This can be extended to multiple environmental variables with additional covariates and corresponding slope terms. However, when plasticity is expressed in response to unknown environmental variables, different approaches are needed.

Double hierarchical generalized linear mixed-effects models (DHGLMs) allow the fitting of random effects on the dispersion terms of models, i.e. the residual variance (Lee and Nelder 2006). Residual variation includes both measurement error and phenotypic plasticity in response to unknown environmental variables (Westneat et al. 2015, Berdal and Dochtermann 2019). Whereas plasticity and buffering in response to known environmental variables can be modeled according to equation 5, DHGLMs enable the detection of plasticity when the variables driving individual responses are unknown or unmeasured.

Including social partners as random effects in a dispersion model captures social buffering in response to unknown environmental variation. A “significant” statistical contribution— however a researcher determines this—of the social partner random term to the dispersion term would then indicate social buffering. Rather than capturing whether social partners increase or decrease the magnitude of a plastic response, social buffering modeled using DGHLMs would indicate an increase or decrease in the absolute magnitude of phenotypic plasticity of focal individuals (Cleasby et al. 2015, Westneat et al. 2015, O’Dea et al. 2022).

Despite this potential, DHGLMs have only rarely been used in evolutionary ecology and, to our knowledge, have not been used for modeling the effects of social partners. Nonetheless, there are an increasing number of options for fitting these models and they can be flexibly and quickly fit in R via C++ coding using the TMB package (Allegue et al. in preparation).

## Conclusion

As we have shown, social buffering—defined as individuals changing the phenotypic plasticity of their social partners—can be readily modeled using mixed-effects models. This modeling is done via the inclusion of social partner identity as a “random slope” term (equation 5). Reliable estimation of variation in social buffering requires large sample sizes (Figures 2 and 3) but these sample sizes are increasingly available to social behavior researchers. This is especially true with automated tracking approaches (Dankert et al. 2009, Dell et al. 2014, Smith and Pinter-Wollman 2021). Importantly, our simulations demonstrate that increasing sampling of social partners and sampling of interactions is more important than increasing sampling of focal individuals. Nonetheless, researchers are encouraged to conduct their own simulations to guide sampling designs (https://osf.io/tg39c).

The general mixed-effects modeling approach we have used here is already frequently used across fields in evolutionary ecology and familiar to many researchers (and reviewers). These approaches are also commonly used for understanding phenotypic plasticity. Modeling social buffering as done here therefore places buffering more clearly into the broader literature on plasticity. The direct connection between biology and statistical approaches illustrated here also facilitates future study of social buffering by providing clear analytical approaches.

Moreover, the phenotypic equation framework encourages researchers to be explicit about their biological hypotheses and assumptions (Allegue et al. 2017, Westneat et al. 2020). In this context, the phenotypic equation framework clarifies the phenotypic effects of social buffering. Specifically, equations 4 and 5 emphasize that social buffering represents an indirect effect of social partners. Indirect effects, defined as how an individual’s phenotype changes the phenotype of a social partner, is most commonly considered in terms of effects on average phenotypes (e.g. equation 4; Bijma 2010) but equally applies to the effects on plasticity captured by equation 5.

Indirect effects and, particularly indirect genetic effects, have been a major focus of research in quantitative genetics. However, to our knowledge, social buffering has not previously been placed in the context of indirect effects. Reconsidering social buffering in this context is potentially important in understanding the evolution of social buffering and the evolution of responses to environmental change. One of the major predictions of indirect genetic effects theory is that these effects can change rates of evolutionary change by changing the amount of genetic variation available for selection to act on (Moore et al. 1997, Santostefano et al. 2024). While theoretically, this can either increase or decrease evolutionary rates, a recent meta-analysis (Santostefano et al. 2024) found that indirect genetic effects, on average, increase the genetic variation on which selection can act. This allows for an accelerated response to selection. If variation in the buffering effects of social partners are underpinned by genetic variation, then plasticity might evolve more rapidly than expected. Whether such genetic variation is common in populations and how this affects the evolution of plasticity is currently unknown and is an important question for future social buffering research.

## Data Availability

All code and data are available at: https://osf.io/t9vxg/

## References Cited

Allegue, H., Y. G. Araya-Ajoy, N. J. Dingemanse, N. A. Dochtermann, L. Z. Garamszegi, S. Nakagawa, D. Reale, H. Schielzeth, and D. F. Westneat. 2017. Statistical Quantification of Individual Differences (SQuID): an educational and statistical tool for understanding multilevel phenotypic data in linear mixed models. Methods in Ecology and Evolution 8:257–267.

Bailey, N. W., L. Marie-Orleach, and A. J. Moore. 2018. Indirect genetic effects in behavioral ecology: does behavior play a special role in evolution? Behavioral Ecology 29:1–11.

Bell, A. M., S. J. Hankson, and K. L. Laskowski. 2009. The repeatability of behaviour: a meta-analysis. Animal Behaviour 77:771–783.

Berdal, M. A., and N. A. Dochtermann. 2019. Adaptive alignment of plasticity with genetic variation and selection. Journal of Heredity 110.

Bijma, P. 2010. Estimating Indirect Genetic Effects: Precision of Estimates and Optimum Designs. Genetics 186:1013–1028.

Brooks, M. E., K. Kristensen, K. J. Van Benthem, A. Magnusson, C. W. Berg, A. Nielsen, H. J. Skaug, M. Machler, and B. M. Bolker. 2017. glmmTMB balances speed and flexibility among packages for zero-inflated generalized linear mixed modeling. The R journal 9:378–400.

Cleasby, I. R., S. Nakagawa, and H. Schielzeth. 2015. Quantifying the predictability of behaviour: statistical approaches for the study of between-individual variation in the within-individual variance. Methods in Ecology and Evolution 6:27–37.

Coe, C. L., D. Franklin, E. R. Smith, and S. Levine. 1982. Hormonal responses accompanying fear and agitation in the squirrel monkey. Physiology & Behavior 29:1051–1057.

Cohen, J. 1988. Statistical power analysis for the behavioral sciences. routledge.

Dankert, H., L. Wang, E. D. Hoopfer, D. J. Anderson, and P. Perona. 2009. Automated monitoring and analysis of social behavior in Drosophila. Nature methods 6:297–303.

Dell, A. I., J. A. Bender, K. Branson, I. D. Couzin, G. G. de Polavieja, L. P. Noldus, A. Pe rez-Escudero, P. Perona, A. D. Straw, and M. Wikelski. 2014. Automated image-based tracking and its application in ecology. Trends in Ecology & Evolution 29:417–428.

Dingemanse, N. J., Y. G. Araya-Ajoy, and D. F. Westneat. 2021. Most published selection gradients are underestimated: Why this is and how to fix it. Evolution 75:806–818.

Dingemanse, N. J., and N. A. Dochtermann. 2013. Quantifying individual variation in behaviour: mixed-effect modelling approaches. Journal of Animal Ecology 82:39–54.

Dingemanse, N. J., N. A. Dochtermann, and S. Nakagawa. 2012. Defining behavioural syndromes and the role of “syndrome” deviation in understanding their evolution. Behavioral Ecology and Sociobiology 66:1543–1548.

Dingemanse, N. J., A. J. Kazem, D. Reale, and J. Wright. 2010. Behavioural reaction norms: animal personality meets individual plasticity. Trends in Ecology & Evolution 25:81–89.

Faber, N. K. M. 1999. Estimating the uncertainty in estimates of root mean square error of prediction: application to determining the size of an adequate test set in multivariate calibration. Chemometrics and Intelligent Laboratory Systems 49:79–89.

Harrison, X. A., L. Donaldson, M. E. Correa-Cano, J. Evans, D. N. Fisher, C. E. Goodwin, B. S. Robinson, D. J. Hodgson, and R. Inger. 2018. A brief introduction to mixed effects modelling and multi-model inference in ecology. Peerj 6:e4794.

Hartman, C. R. A., G. S. Wilkinson, I. Razik, I. M. Hamilton, E. A. Hobson, and G. G. Carter. 2024. Hierarchically embedded scales of movement shape the social networks of vampire bats. Proceedings of the Royal Society B 291:20232880.

IPCC. 2023. Climate Change 2023: Synthesis Report. Geneva, Switzerland.

Kikusui, T., J. T. Winslow, and Y. Mori. 2006. Social buffering: relief from stress and anxiety. Philosophical Transactions of the Royal Society B: Biological Sciences 361:2215–2228.

Kruuk, L. E. B. 2004. Estimating genetic parameters in natural populations using the ‘animal model’. Philosophical Transactions of the Royal Society of London Series B-Biological Sciences 359:873–890.

Lawson, C. R., Y. Vindenes, L. Bailey, and M. van de Pol. 2015. Environmental variation and population responses to global change. Ecology Letters 18:724–736.

Lee, Y., and J. A. Nelder. 2006. Double hierarchical generalized linear models (with discussion). Journal of the Royal Statistical Society Series C: Applied Statistics 55:139–185.

MacLeod, K. J., C. Monestier, M. C. Ferrari, K. E. McGhee, M. J. Sheriff, and A. M. Bell. 2022. Predator-induced transgenerational plasticity in animals: a meta-analysis. Oecologia 200:371–383.

Martin, J. S., and A. V. Jaeggi. 2022. Social animal models for quantifying plasticity, assortment, and selection on interacting phenotypes. Journal of Evolutionary Biology 35:520–538.

Moore, A. J., E. D. Brodie, and J. B. Wolf. 1997. Interacting phenotypes and the evolutionary process .1. Direct and indirect genetic effects of social interactions. Evolution 51:1352–1362.

Noble, D. W., V. Stenhouse, and L. E. Schwanz. 2018. Developmental temperatures and phenotypic plasticity in reptiles: A systematic review and meta-analysis. Biological Reviews 93:72–97.

Nussey, D. H., E. Postma, P. Gienapp, and M. E. Visser. 2005. Selection on heritable phenotypic plasticity in a wild bird population. Science 310:304–306.

Nussey, D. H., A. J. Wilson, and J. E. Brommer. 2007. The evolutionary ecology of individual phenotypic plasticity in wild populations. Journal of Evolutionary Biology 20:831–844.

O’Dea, R. E., D. W. Noble, and S. Nakagawa. 2022. Unifying individual differences in personality, predictability and plasticity: a practical guide. Methods in Ecology and Evolution 13:278–293.

Ogino, M., A. A. Maldonado-Chaparro, L. M. Aplin, and D. R. Farine. 2023. Group-level differences in social network structure remain repeatable after accounting for environmental drivers. Royal Society Open Science 10:230340.

Ponzi, E., L. F. Keller, T. Bonnet, and S. Muff. 2018. Heritability, selection, and the response to selection in the presence of phenotypic measurement error: Effects, cures, and the role of repeated measurements. Evolution 72:1992–2004.

Santostefano, F., M. Moiron, A. Sanchez-Tojar, and D. N. Fisher. 2024. Indirect genetic effects increase the heritable variation available to selection and are largest for behaviors: a meta-analysis. Evolution Letters: qrae051.

Smith, J. E., D. A. Gamboa, J. M. Spencer, S. J. Travenick, C. A. Ortiz, R. D. Hunter, and A. Sih. 2018. Split between two worlds: automated sensing reveals links between above- and belowground social networks in a free-living mammal. Philosophical Transactions of the Royal Society B: Biological Sciences 373:20170249.

Smith, J. E., and N. Pinter-Wollman. 2021. Observing the unwatchable: Integrating automated sensing, naturalistic observations and animal social network analysis in the age of big data. Journal of Animal Ecology 90:62–75.

Stamp, M. A., and J. D. Hadfield. 2020. The relative importance of plasticity versus genetic differentiation in explaining between population differences; a meta-analysis. Ecology Letters 23:1432–1441.

Westneat, D. F., Y. G. Araya-Ajoy, H. Allegue, B. Class, N. Dingemanse, N. A. Dochtermann, L. Z. Garamszegi, J. G. Martin, S. Nakagawa, and D. Reale. 2020. Collision between biological process and statistical analysis revealed by mean centring. Journal of Animal Ecology 89:2813–2824.

Westneat, D. F., J. Wright, and N. J. Dingemanse. 2015. The biology hidden inside residual within-individual phenotypic variation. Biological Reviews 90:729–743.

Wolf, J. B., E. D. Brodie, J. M. Cheverud, A. J. Moore, and M. J. Wade. 1998. Evolutionary consequences of indirect genetic effects. Trends in Ecology & Evolution 13:64–69.

Wolf, J. B., E. D. Brodie, and A. J. Moore. 1999. Interacting phenotypes and the evolutionary process. II. Selection resulting from social interactions. American Naturalist 153:254–266.

Young, C., B. Majolo, M. Heistermann, O. Schu lke, and J. Ostner. 2014. Responses to social and environmental stress are attenuated by strong male bonds in wild macaques. Proceedings of the National Academy of Sciences 111:18195–18200.

